# Breast adipose tissue-derived extracellular vesicles from women with obesity stimulate mitochondrial-induced dysregulated tumor cell metabolism

**DOI:** 10.1101/2023.02.08.527715

**Authors:** Shuchen Liu, Alberto Benito-Martin, Fanny A. Pelissier Vatter, Sarah Z. Hanif, Catherine Liu, Priya Bhardwaj, Praveen Sethupathy, Alaa R. Farghli, Phoebe Piloco, Paul Paik, Malik Mushannen, David M. Otterburn, Leslie Cohen, Rohan Bareja, Jan Krumsiek, Leona Cohen-Gould, Samuel Calto, Jason A. Spector, Olivier Elemento, David Lyden, Kristy A. Brown

## Abstract

Breast adipose tissue is an important contributor to the obesity-breast cancer link. Dysregulated cell metabolism is now an accepted hallmark of cancer. Extracellular vesicles (EVs) are nanosized particles containing selective cargo, such as miRNAs, that act locally or circulate to distant sites to modulate target cell functions. Here, we found that long-term education of breast cancer cells (MCF7, T47D) with EVs from breast adipose tissue of women who are overweight or obese (O-EVs) leads to sustained increased proliferative potential. RNA-Seq of O-EV-educated cells demonstrates increased expression of genes, such as ATP synthase and NADH: ubiquinone oxidoreductase, involved in oxidative phosphorylation. O-EVs increase respiratory complex protein expression, mitochondrial density, and mitochondrial respiration in tumor cells. Mitochondrial complex I inhibitor, metformin, reverses O-EV-induced cell proliferation. Several miRNAs, miR-155-5p, miR-10a-3p, and miR-30a-3p, which promote mitochondrial respiration and proliferation, are enriched in O-EVs relative to EVs from lean women. O-EV-induced proliferation and mitochondrial activity are associated with stimulation of the Akt/mTOR/P70S6K pathway, and are reversed upon silencing of P70S6K. This study reveals a new facet of the obesity-breast cancer link with human breast adipose tissue-derived EVs causing the metabolic reprogramming of ER+ breast cancer cells.

## Introduction

Breast cancer is one of the most common cancers in women, with 2.3 million new cases diagnosed each year ^1^. Excess adiposity, as seen in people with overweight or obesity (body-mass index [BMI] ≥ 25kg/m^2^), is recognized as an important risk factor for at least 13 different cancers, including postmenopausal, hormone receptor-positive breast cancer ^2^. Pathophysiological aspects of obesity include alterations in metabolic health that occur because of adipose tissue dysfunction, which has also been proposed to be a key driver of tumor development and progression ^3^. Extracellular vesicles (EVs) are key players in the tumor microenvironment, allowing for direct communication between different cell types and having been shown to play an important role in metastatic dissemination and cancer progression ^4–6^. All cells, including adipocytes, mesenchymal stem cells and immune cells within the adipose tissue microenvironment, constitutively release EVs into the extracellular space ^7^. Adipose tissue-derived EVs have been shown to communicate directly with macrophages ^8–10^, monocytes ^11^, and endothelial cells ^12^. Moreover, miRNAs contained in adipose tissue-derived EVs have been shown to act systemically to influence distal organs ^13,14^. The adipose tissue-tumor interplay has been extensively studied ^15^, but a role for EVs has only recently been proposed. Lazar *et al*. showed that adipocytes secrete EVs in abundance, which are then taken up by melanoma cells, leading to increased cell migration and invasion ^16^. A reciprocal communication between breast cancer cells and adipose tissue has also been described, including increased lipolysis and release of fatty acids from adipocytes in response to tumor cells and support of malignant progression and resistance to chemotherapeutic agents ^17–22^. Recently, it was demonstrated that EVs isolated from conditioned media of *in vitro*-differentiated adipocytes induced phenotypic changes consistent with epithelial-to-mesenchymal transition (EMT) ^23^. Little is known of the impact of obesity on the content and effect of EVs from human adipose tissue in driving breast cancer development and progression.

In this study, we characterized EVs released by human breast adipose tissue obtained from reduction mammoplasty, including how cargo is altered with obesity, and examined effects of long-term education of these EVs on the proliferation of estrogen receptor + (ER+) breast cancer cells. We also demonstrate a shift in cell metabolism towards oxidative phosphorylation that supports increased rates of cell proliferation, pointing to a role for EVs in metabolic reprogramming of their recipient cells.

## Materials and Methods

### Human breast adipose tissue

Human breast adipose tissue was obtained from patients undergoing reduction mammoplasty at Weill Cornell Medicine under IRB-approved protocols (protocol #20-01021391 and #1510016712). All patients provided written informed consent for collection, storage, distribution of samples and data analysis for research studies. The clinical information of all patients was retrieved from their electronic medical records. BMI was recorded prior to surgery. Samples were separated into two groups: lean (BMI < 25 kg/m^2^) and overweight/obese (BMI ≥ 25 kg/m^2^).

### EV isolation and characterization

Human breast adipose tissue was incubated in mammary epithelial cell growth base medium (MEBM) (Lonza #cc-3151) supplemented with 0.5% bovine serum albumin (BSA) and depleted of EVs by ultra-centrifugation (100,000g, 70 minutes). Ten 1cm^3^ pieces of breast adipose tissue were incubated for 24 hours at 37°C (10 blocks/10cm dish) in a humidified atmosphere with 5% CO_2_, and the supernatant/conditioned media was collected. EV isolation was performed as described previously ^24^. Briefly, conditioned media was sequentially centrifuged at 300xg for 5 minutes, 500xg for 10 minutes, 3,000xg for 20 minutes and 12,000xg for 20 minutes. EVs were then pellets from the supernatant via centrifugation at 100,000xg for 70 minutes, followed by an additional centrifugation at 100,000xg. All steps were performed at 4 °C. EVs were resuspended in PBS and stored at −20 °C. Collected EVs were then characterized using BCA (Pierce Biotechnology) to measure protein concentration and the NanoSight NS500 (Malvern Panalytical) system and Nanoparticle Tracking Analysis (NTA) software 2.3 to characterize EV diameter and concentration.

### Transmission electron microscopy (TEM)

For TEM of EVs, samples were fixed with 2% paraformaldehyde were placed on glow discharged formvar-carbon coated nickel grids. Grids were blocked for 10 min with 1% fish skin gelatin (Sigma-Aldrich). The grids were then washed with PBS and fixed in 1% glutaraldehyde for 5 min and then embedded in a mixture of 3% uranyl acetate and 2% methylcellulose. Stained grids were examined under Philips CM-12 electron microscope and photographed with a digital camera.

For TEM of long-term educated cells, cells were plated in a 6-well plate and imaged when they reached ^~^80% confluency, as described previously ^25^. Briefly, cells were fixed with fixation solution (4% paraformaldehyde, 2.5% glutaraldehyde, 0.02% picric acid in 0.1M sodium cacodylate buffer, pH 7.3), stained with uranyl acetate, and dehydrated with a graded ethanol series. After dehydration, samples were covered by a layer of fresh resin, and embedding molds were inserted into the resin. After polymerization, samples were cut at 200nm for screening by light microscopy and then at 70nm to be mounted on grids for TEM under a JEOL JSM 1400 (JEOL, USA) electron microscope. The camera used is a Veleta, 2K x 2K CCD (EMSIS, GmbH, Muenster). Images were taken under 8000X magnification, and each field included extracellular space, cytoplasmic area and nucleus. We randomly selected two sections for each case and 6 fields were randomly imaged for each section. Mitochondrial area and cytoplasmic area were identified by image J. Mitochondrial density was evaluated by counting the number of mitochondria per μm^2^ of cytoplasmic area.

### Proteomic analysis

EVs from 48 cases were analyzed for protein content. Samples were reduced and alkylated with 5mM DTT and 14mM iodoacetamide, respectively, and then trypsinized overnight at 37 C. Digests were then lyophilized and desalted by micro-C18 columns. Proteomic analysis was performed as described previously ^26^. Briefly, samples were analyzed using a Thermo Fisher Scientific EASY-nLC 1000 coupled online to a Fusion Lumos mass spectrometer (Thermo Fisher Scientific) with peptide separation performed using a 75 μm x 15 cm chromatography column (ReproSil-Pur C18-AQ, 3 μm, Dr. Maisch GmbH, German) packed in-house. Full MS scans were acquired in the Orbitrap mass analyzer over a range of 300-1500 m/z with resolution 60,000. MS/MS scans were acquired in the Orbitrap mass analyzer with resolution 15,000. The raw files were processed using the MaxQuant computational proteomics platform version 1.5.5.1 (Max Planck Institute, Munich, Germany) for protein identification. The fragmentation spectra were used to search the UniProt human protein database (downloaded on 09/21/2017). Both peptide and protein identifications were filtered at 1% false discovery rate based on decoy search using a database with the protein sequences reversed.

### Cell culture

Human breast cancer cells MCF7 (Cat# HTB-22) and T47D (Cat# HTB-133) were obtained from the ATCC. Murine breast cancer cells EO771 (#94A001) were purchased from CH3 BioSystems. MCF7 and EO771 cells were cultured in DMEM (Gibco #11995) supplemented with 10% fetal bovine serum (FBS) (Gibco 26140079) depleted of EVs by ultra-centrifugation (100,000g, 70 minutes) and 1% penicillin/streptomycin at 37°C in a humidified atmosphere with 5% CO2. T47D cells were cultured in RPMI Medium 1640 (Gibco 11875) supplemented with 10% fetal bovine serum (FBS) depleted of EVs and 1% penicillin/streptomycin. All cell lines were tested for mycoplasma contamination using the Universal Mycoplasma Detection Kit (ATCC 30-1012K).

### Single-dose treatment with EVs

MCF7 cells were seeded in complete media at a density of 10,000 cells/well of the E-Plate of Agilent ACEA xCELLigence, see method of xCELLigence real-time cell assay (RTCA) system for more details. After 24 hours, cells were treated with media containing EVs (1 μg/ml) from adipose tissue of patients with overweight/obesity (O-EVs), with a healthy weight (lean; L-EVs) or vehicle control (PBS; same volume). Impedance measurements were taken for 90 hours.

### Long-term education (LTE) with EVs or miRNA mimics

MCF7 and T47D cells were seeded at 40,000 cells/well in a 6-well plate, and treated with L-EVs, O-EVs or vehicle control (PBS; same volume) on day 1, 4 and 7. The culture media was refreshed with each EV treatment. On day 8, cells were harvested for further experiments.

Long-term education of breast cancer cell lines with miRNA mimics was performed using a similar strategy. On day 1, 4 and 7, culture medium was replaced with base medium 1 hour before transfection and cells were transfected with miRNA mimics (Qiagen), including hsa-let-7a-5p (UGAGGUAGUAGGUUGUAUAGUU), hsa-miR-155-5p (UUAAUGCUAAUCGUGAUAGGGGU), hsa-miR-10a-3p (CAAAUUCGUAUCUAGGGGAAUA), hsa-miR-29b-3p (UAGCACCAUUUGAAAUCAGUGUU), hsa-miR-30a-3p (CUUUCAGUCGGAUGUUUGCAGC), hsa-miR-30b-5p (UGUAAACAUCCUACACUCAGCU) and negative control (UCACCGGGUGUAAAUCAGCUUG), using Lipofectamine RNAiMAX (Thermo-Fisher Scientific). The base medium containing miRNA mimics – lipo complexes was replaced with normal culture media after 6 hours. On day 8, cells were harvested for further experiments.

### xCELLigence real-time cell assay (RTCA) system for cell proliferation

The Agilent ACEA xCELLigence RTCA DP Real-Time Cell Analyzer (Agilent, 3X16) was used to monitor cell proliferation after long-term education. Educated MCF7 or T47D cells were seeded at 10,000 cells/well of the E-Plate in a volume of 100 μL culture media and cell index (CI) was monitored over a 90-hour period. ΔCI over time was used to assess differences in cell proliferation rates.

For experiments examining the effect of metformin treatment, cells were seeded in a RTCA E-Plate as described above. After 24 hours, the medium was replaced with medium containing 0.5mM metformin hydrochloride (#PHR1084, Sigma), an inhibitor of mitochondrial ATP synthase and oxidative phosphorylation. CI was monitored over time and proliferation rate measured as above.

### Murine experimental model

All animal experiments were performed at Weill Cornell Medicine in accordance with IACUC protocol #2018-0058. C57BL/6J mice were purchased from JAX laboratories.

Four-week-old female mice were ovariectomized, allowed to recover for 1 week, and randomized to two groups (5 mice per group): 10 kcal% low fat diet (LFD, Research Diets, 12450Bi) or 60 kcal% high fat diet (HFD, Research Diets, D12492i) for ten weeks (Fig. 2G). Body weights were measured every week after the 3^rd^ week on diet. In the 11^th^ week, mice were euthanized to collect mammary fat pads (MFP). All MFPs were collected and pooled based on diet to generate conditioned media from which EVs were isolated (as in EV preparation section). EO771, a murine mammary cancer cell line syngeneic to C57BL/6 mice ^27^, was long-term educated with EVs generated from HFD or LFD groups, as described above.

**Figure 1.**
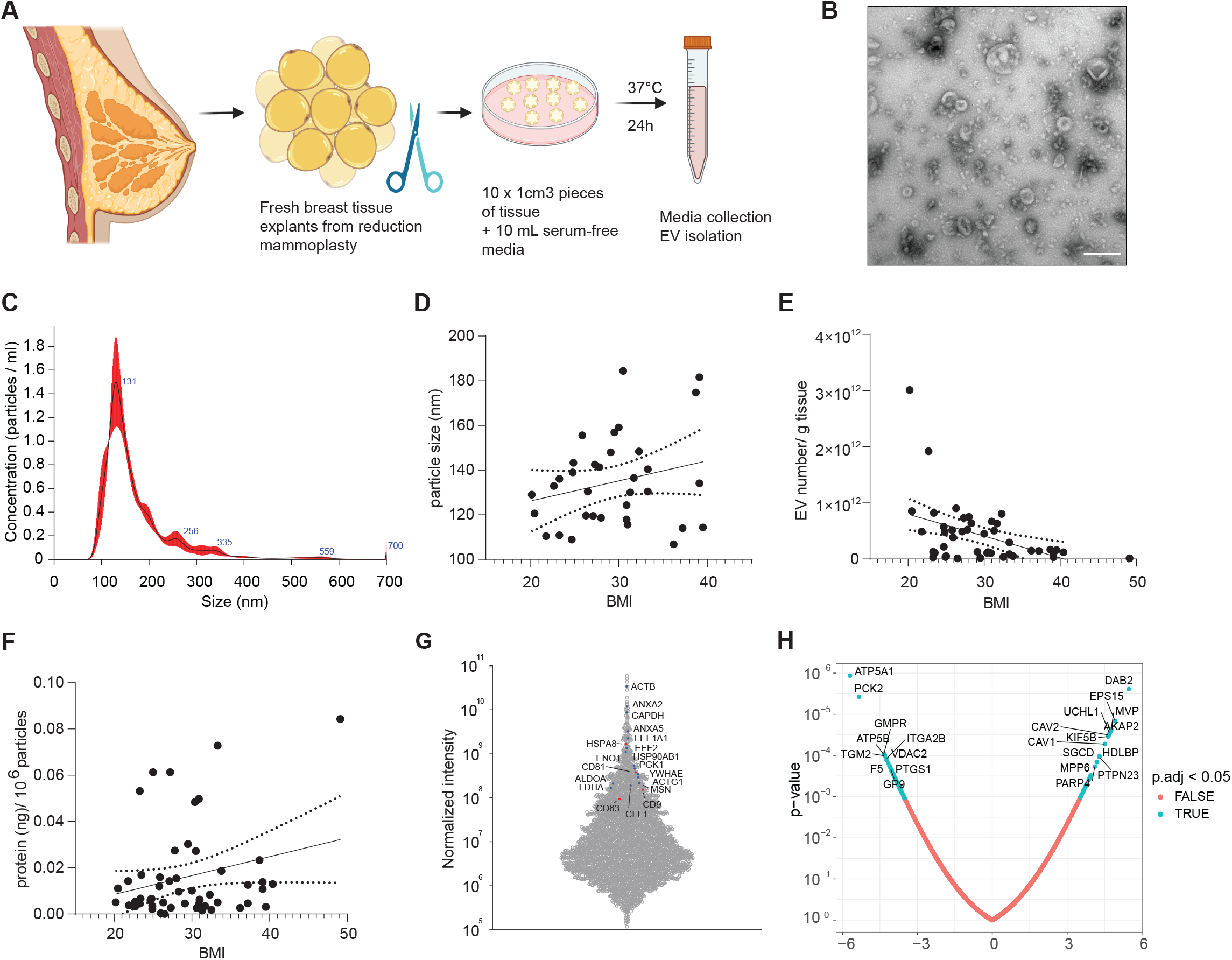
Characterization of EVs derived from human breast adipose tissue. A) Flow diagram of generation of conditioned media from breast adipose tissue. B) Representative TEM image of isolated EVs. Scale bar, 200nm. C) Nanoparticle Tracking Analysis representative graph of collected EVs showing size and relative abundance. (D-F) Linear regression analysis of BMI and (D) particle size (nm), (E) EV number/g tissue, (F) protein (ng)/10^6^ particles. G) Proteomic analysis of EVs from 48 cases indicating relative abundance of known exosome markers. Commonly used EV markers are depicted in red and additional EV markers identified using ExoCarta are depicted in blue. H) Volcano plot reflecting proteins with significant correlations with BMI.

**Figure 2.**
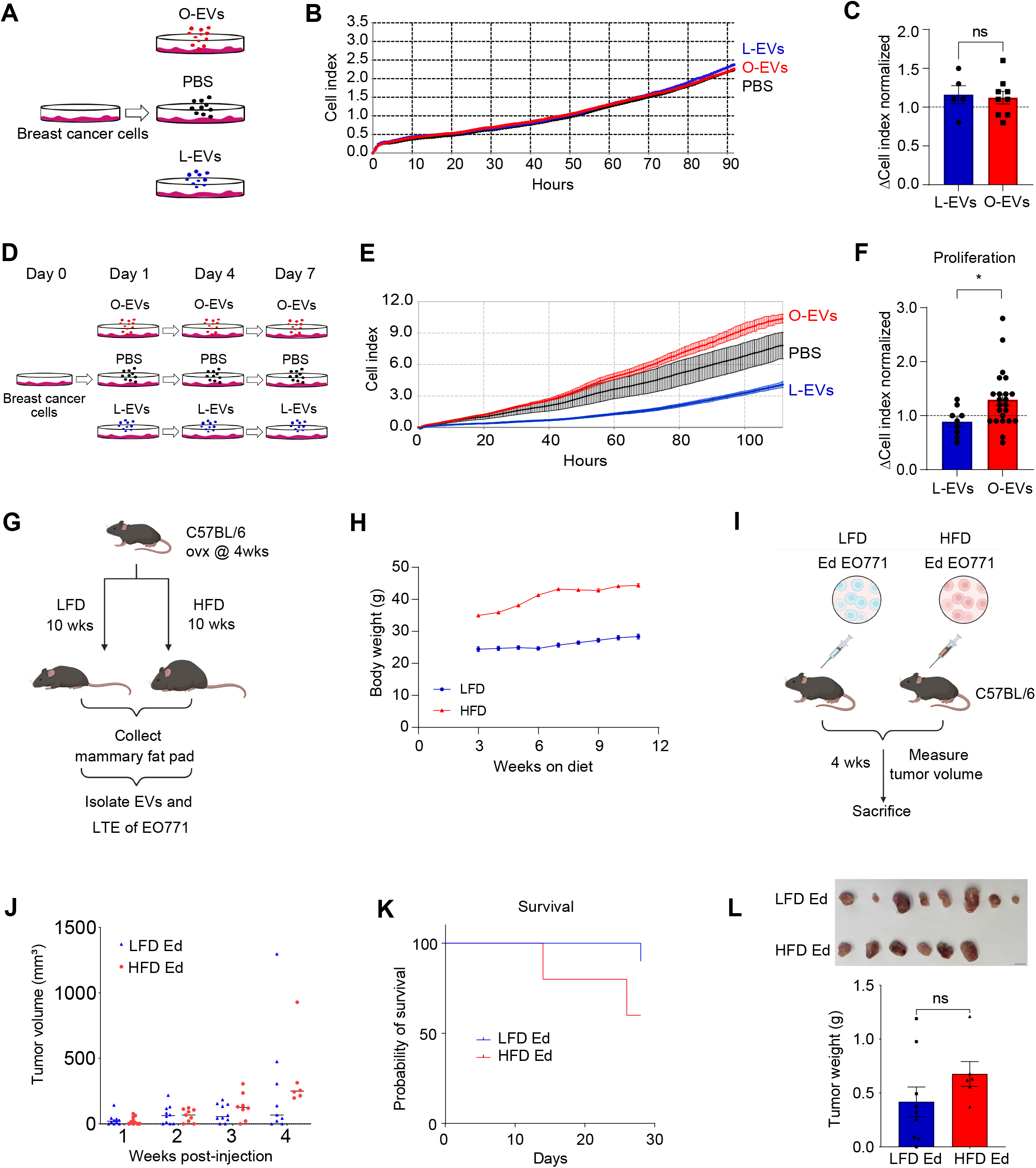
Adipose tissue derived EVs from women with overweight/obesity promote proliferation of breast cancer cells *in vitro* and *in vivo*. A) Flow diagram of single-dose treatment of breast cancer cells with EVs and PBS (control). B) Representative xCELLigence graph showing the effect of single-dose treatment with O-EVs, L-EVs or PBS on MCF7 cell proliferation. C) Quantification of relative proliferation rates of MCF7 cells educated with O-EVs (n=9) vs L-EVs (n=5). Proliferation rate was determined by measuring the slope of the cell index within a 20-hour time frame within the linear phase of cell division. Proliferation rate of L-EV- and O-EV-educated cells was normalized to their controls. D) Experimental schematic of long-term education of breast cancer cells with isolated EVs or PBS (control). E) Representative xCELLigence image demonstrating the effect of long-term education with O-EVs or L-EVs on MCF7 cells proliferation. F) Comparison of relative proliferation of MCF7 cells educated with O-EVs (n=23) vs L-EVs (n=9). G) Flow diagram of generation of EVs from low-fat diet (LFD) or high-fat diet (HFD) fed mice and long-term education of EO771 cells. H) Weekly body weight measurements of mice fed LFD or HFD for 12 weeks. I) Schematic of allografting of C57Bl/6 mice with HFD- or LFD-EV long-term educated E0771 cells via mammary fat pad injection. J) Weekly tumor growth of LFD-EV-educated (LFD Ed) and HFD-EV-educated (HFD Ed) E0771 following mammary fat pad injection. K) Survival analysis of LFD Ed and HFD Ed allografted mice over 28 days. L) Picture of isolated tumors after euthanasia and tumor weight comparison, p=0.0879. Scale bar, 1 cm. Bars and error bars represent means ± SEM; statistically significant by unpaired t-test with Welch’s correction (C) or Mann-Whitney test (F, L), (ns P≥ 0.05, *P <0.05).

Eight-week-old C57BL/6 female mice were randomized to two groups (10 mice per group), receiving a mammary fat pad injection (5×10^5 cells per mouse) of EO771 cells educated with EVs from LFD Ed group or HFD Ed group, respectively. Tumors were identified by palpation and measured with digital electronic caliper (Finescience, 30087-00) every week after injection. After 4 weeks, the mice were euthanized to isolate tumors.

### RNA preparation and RNA-Seq analysis

Cells were lysed using Qiazol lysis reagent (Qiagen, 79306), and total RNA was extracted from samples by Trizol lysis and RNeasy Mini Kit (Qiagen) following the manufacturer’s protocols. Samples were submitted for RNA-seq at the Genomics Resources Core Facility (GRCF, Weill Cornell Medicine). Sequencing libraries were constructed at the GRCF following the Illumina TruSeq Stranded mRNA library preparation protocol (Poly-A selection and Stranded RNA-Seq). The libraries were sequenced with paired-end 50 bp on the Illumina HiSeq4000 sequencing platform. All reads were independently aligned with STAR_2.4.0f1 ^28^ for sequence alignment against the human genome sequence build hg19, downloaded via the UCSC genome browser and SAMTOOLS v0.1.19 ^29^ for sorting and indexing reads. Cufflinks (2.0.2) ^30^ was used to estimate the expression values (FPKMS), and GENCODE v19 ^31^ GTF file for annotation. The gene counts from htseq-count and DESeq2 Bioconductor package ^32^ were used to identify DEGs. All DEGs with *P* value <0.05 were uploaded to IPA for data visualization and analysis.

### miRNA analysis

Total RNA was extracted from EVs using the mirVana miRNA isolation kit (Thermo Fisher Scientific) as recommended by the manufacturer. RNA purity was quantified with the Nanodrop 2000 instrument (Thermo Fisher Scientific), and RNA integrity was quantified with the Agilent 2100 Bioanylyzer Inmstrument (Aglient Technologies). Libraries were generated using the CleanTag Small RNA Library Prep Kit (TriLink Biotechnologies). Sequencing was performed on the HiSeq3000 platform (Illumina) at the Genome Sequencing Facility of the Greehey Children’s Cancer Research Institute (University of Texas Health Science Center). Read quality was assessed using FastQC. Trimming, mapping and quantification was performed using miRquant 2.0.

### Seahorse assay

Mitochondrial respiration was evaluated using the Seahorse XF Cell Mito Stress Test Kit (Agilent 103015-100) with the Seahorse XFe96 Analyzer (Agilent), following manufacturer’s Mito Stress Test protocol. Briefly, 20,000 cells/well were plated in a 96-well Seahorse assay plate and the cartridge was hydrated one day before assay. On the day of assay, the culture medium in the assay plate was replaced with XF assay medium modified DMEM (102365-100, Agilent) supplemented with 1 mM Pyruvate, 2 mM Glutamine and 10mM glucose, and the cells were incubated for one hour at 37 °C in a non-CO2 incubator. Mitochondrial complex inhibitors were loaded sequentially into the ports on sensor cartridge (Oligomycin 1.5 μM, FCCP 0.5 μM and 0.5 μM Rotenone/antimycin A for MCF7 cells; Oligomycin 1.5 μM, FCCP 1 μM and 1 μM Rotenone/antimycin A for T47D cells). OCR (oxygen consumption rate) and ECAR (extracellular acidification rate) were measured three times in each phase to reflect basal respiration, mitochondrial related ATP production and maximal respiratory capacity.

### Western blot analysis

Protein extraction was performed as previously described ^33^. BCA (Pierce Biotechnology) was used to determine protein concentration. Cell extracts were denatured in buffer containing DTT, run on NuPAGE 4-12% Bis-Tris protein gels (Thermo-Fisher Scientific; 50 μg protein per lane) and transferred to nitrocellulose membranes. Membranes were blocked with 5% nonfat dry milk (Bio-Rad, #1706404) for 1 hour in the room temperature (RT) and incubated with primary antibodies in 4 °C overnight. The following primary antibodies were used: phospho-Akt (Ser473) (Cell Signaling #4060), Akt (Cell Signaling #9272), phospho-p70 S6 Kinase (Thr389) (Cell Signaling #9234), p70 S6 Kinase (Cell Signaling #2708), phospho-4EBP1 (Ser65) (Cell Signaling #9451), 4EBP1 (Cell Signaling #9644), OxPhos Human WB Antibody Cocktail (Thermo-Fisher Scientific #45-8199) and β-actin (Sigma-Aldrich, A3854). The following secondary antibodies were used: Anti-rabbit IgG HRP-linked Antibody (Cell Signaling #7074) and Anti-mouse IgG HRP-linked Antibody (Cell Signaling #7076). For phospho-protein and total protein detection, membranes were probed with phospho-protein antibody first, and stripped it with Restore PLUS western blot stripping buffer (Thermo-Fisher Scientific, #46430) for 15 mins. Due to limited material, some membranes were cut to allow probing of multiple targets with distinct molecular weights from a single nitrocellulose membrane. Western lightning plus-ECL (Thermo-Fisher Scientific) and ImageLab software (BioRad) were used for band detection.

### Immunofluorescent staining, imaging, and image analysis

Cells were plated at 20,000 cells/well in a 96-well assay black plate (Corning #3603). On the following day, culture medium was replaced with culture medium containing 100 nM MitoTracker Green FM (MTG) (Thermo-Fisher Scientific M7514) and 20 μM Hoechst. The plate was incubated at 37°C for 30 minutes and imaged at 20x magnification immediately using confocal microscopy (Zeiss LSM880). Imaris x64© software (version 9.7.2, OXFORD instruments) was used for image analysis; all settings and parameters were standardized across the images. Statistical analysis was exported as “cell cytoplasm intensity mean” of green fluorescent signal to reflect mitochondrial mass of each cell. The intensity mean value per cell for each sample were then normalized to the average intensity mean of all cells in the respective control sample.

### siRNA targeting P70S6K

Gene silencing of P70S6K in T47D cells was carried out using siRNA. Cells at 60-70% confluence were incubated in Opti-MEM (Gibco #31985-070) for 1 hour and then transfected with siRNAs (Qiagen, Cat. No. 1027417) using lipofectamine RNAiMAX (Thermo-Fisher Scientific #13778) following the manufacturer’s protocol. The following siRNAs were used: Hs_RPS6KB1_1 (also known as siP70S6K1_1): AAGAGGGTTCTTTATGTTATA or control siRNA AATTCTCCGAACGTGTCACGT. Media was changed to complete media after 6hrs. Cells were collected after 24 hours. The efficiency of knockdown of P70S6K was assessed by Western blot. The effects of P70S6K on cell proliferation and mitochondrial respiration were measured by xCELLigence and Seahorse assays, as above.

### Statistical analysis

All experiments were performed at least two times, in triplicate. Statistical analyses and graphical presentation were carried out using GraphPad Prism 8.0. All data were reported as mean ± SEM. Student *t* test, one sample t test or one-way ANOVA were used to assess statistical significance (as shown in the figure legends). A p value <0.05 was considered statistically significant (*, p<0.05; **, p<0.005; ***, p<0.0005; ****, p<0.0001; n.s., not significant).

## Results

### Characterization of breast adipose tissue-derived EVs

Breast adipose tissue-derived EVs were isolated from conditioned media (n=52) via ultracentrifugation (Fig. 1A) and displayed a characteristic cup-like shape on TEM (Fig. 1B). The size of EVs for a subset of these cases was evaluated and ranged between 50-300 nm, with approximately 80% of particles having a diameter of between 100-200 nm (Fig. 1C). There was no significant difference in median particle size across BMI of the subjects (Fig. 1D, *r* = 0.2432, p = 0.1657), although the number of particles per gram (g) of breast adipose tissue was inversely associated with the BMI of women from which they were obtained (Fig. 1E; *r* = −0.4443, p = 0.0025). There is a trend for EV protein content being positively associated with BMI (Fig. 1F; *r* = 0.2471, p = 0.0774). Proteomic analysis of EVs from 48 surgical cases demonstrates high abundance of well-characterized exosome-specific markers, including HSPA8, CD81, CD9 and CD63, as well as other established exosome markers based on ExoCarta (Fig. 1G). The abundance of 74 proteins was significantly correlated with BMI, with the top 10 positively associated proteins being A-Kinase Anchoring Protein 2 (AKAP2), Ubiquitin C-terminal Hydrolase L1 (UCHL1), Defective In Cullin Neddylation 1 Domain Containing 3 (DCUN1D3), Epidermal Growth Factor Receptor Pathway Substrate 15 (EPS15), Protein Tyrosine Phosphatase Non-receptor type 23 (PTPN23), MAGUK p55 subfamily member 6 (MPP6), Canine Adenovirus type 2 (CAV2), Sorbin and SH3 domain-containing protein 1 (SORBS1), Poly(ADP-ribose) Polymerase family member 4 (PARP4), Protein Geranylgeranyltransferase Type I, beta subunit (PGGT1B), while the top 10 negatively associated proteins are Coagulation factor V (F5), Serpin family A member 1 (SERPINA1), Latent Transforming Growth Factor beta Binding Protein 1 (LTBP1), Adducin 2 (ADD2), Quiescin Sulfhydryl Oxidase 1 (QSOX1), Hemoglobin Subunit Theta 1 (HBQ1), Serpin family A member 4 (SERPINA4), Phosphoenolpyruvate Carboxykinase 2 (PCK2), Carcinoembryonic Antigen-related Cell Adhesion Molecule 8 (CEACAM8), Protein S (PROS1) (Fig. 1H). The top 30 EV proteins positively associated with BMI and the top 30 EV proteins negatively associated with BMI are presented in Table 1.

**Table 1.**
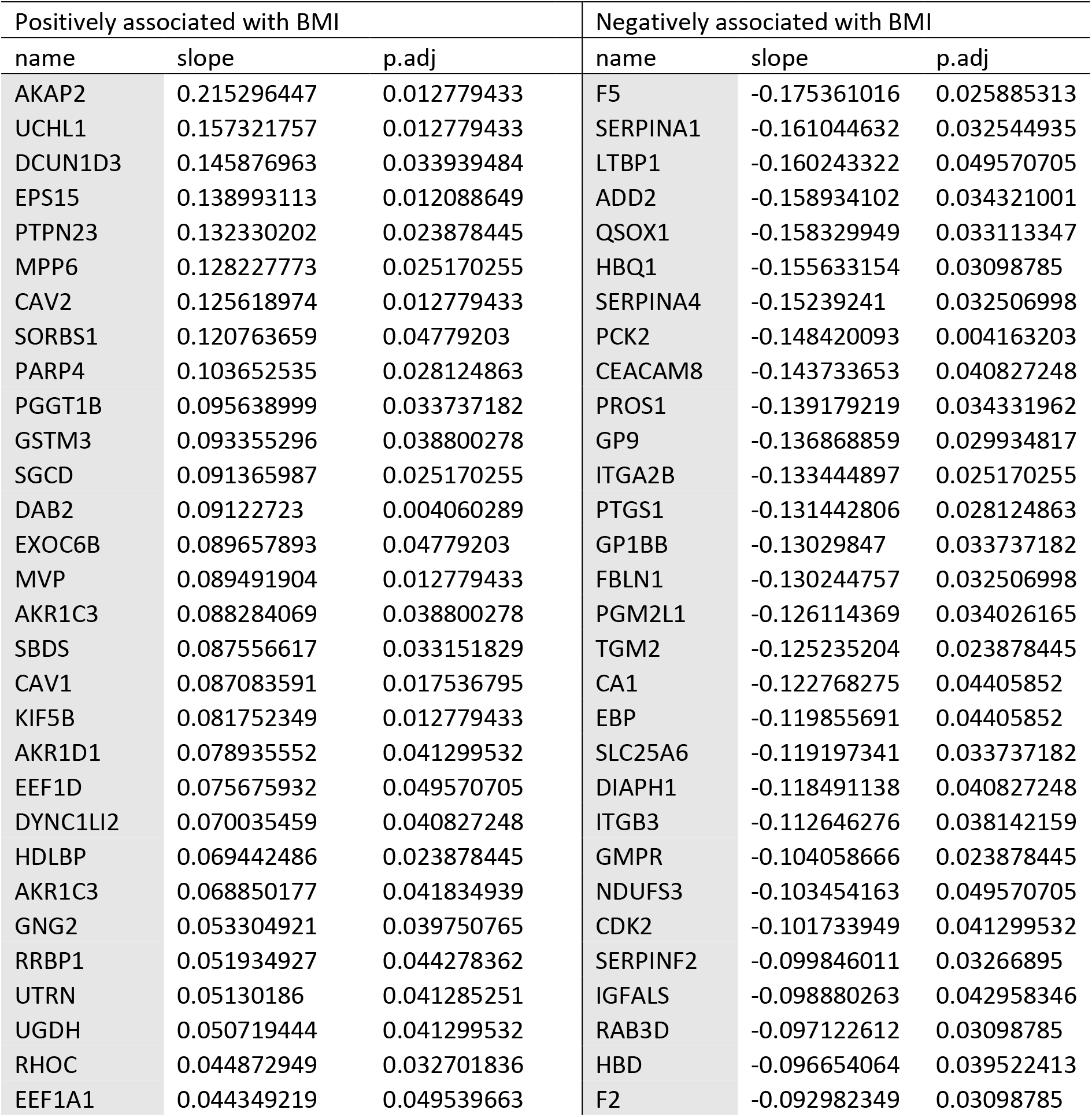
Top 30 EV proteins significantly associated with BMI

| Positively associated with BMI |  |  | Negatively associated with BMI |  |  |
| --- | --- | --- | --- | --- | --- |
| name | slope | p.adj | name | slope | p.adj |
| AKAP2 | 0.215296447 | 0.012779433 | F5 | -0.175361016 | 0.025885313 |
| UCHL1 | 0.157321757 | 0.012779433 | SERPINA1 | -0.161044632 | 0.032544935 |
| DCUN1D3 | 0.145876963 | 0.033939484 | LTBP1 | -0.160243322 | 0.049570705 |
| EPS15 | 0.138993113 | 0.012088649 | ADD2 | -0.158934102 | 0.034321001 |
| PTPN23 | 0.132330202 | 0.023878445 | QSOX1 | -0.158329949 | 0.033113347 |
| MPP6 | 0.128227773 | 0.025170255 | HBQ1 | -0.155633154 | 0.03098785 |
| CAV2 | 0.125618974 | 0.012779433 | SERPINA4 | -0.15239241 | 0.032506998 |
| SORBS1 | 0.120763659 | 0.04779203 | PCK2 | -0.148420093 | 0.004163203 |
| PARP4 | 0.103652535 | 0.028124863 | CEACAM8 | -0.143733653 | 0.040827248 |
| PGGT1B | 0.095638999 | 0.033737182 | PROS1 | -0.139179219 | 0.034331962 |
| GSTM3 | 0.093355296 | 0.038800278 | GP9 | -0.136868859 | 0.029934817 |
| SGCD | 0.091365987 | 0.025170255 | ITGA2B | -0.133444897 | 0.025170255 |
| DAB2 | 0.09122723 | 0.004060289 | PTGS1 | -0.131442806 | 0.028124863 |
| EXOC6B | 0.089657893 | 0.04779203 | GP1BB | -0.13029847 | 0.033737182 |
| MVP | 0.089491904 | 0.012779433 | FBLN1 | -0.130244757 | 0.032506998 |
| AKR1C3 | 0.088284069 | 0.038800278 | PGM2L1 | -0.126114369 | 0.034026165 |
| SBDS | 0.087556617 | 0.033151829 | TGM2 | -0.125235204 | 0.023878445 |
| CAV1 | 0.087083591 | 0.017536795 | CA1 | -0.122768275 | 0.04405852 |
| KIF5B | 0.081752349 | 0.012779433 | EBP | -0.119855691 | 0.04405852 |
| AKR1D1 | 0.078935552 | 0.041299532 | SLC25A6 | -0.119197341 | 0.033737182 |
| EEF1D | 0.075675932 | 0.049570705 | DIAPH1 | -0.118491138 | 0.040827248 |
| DYNC1LI2 | 0.070035459 | 0.040827248 | ITGB3 | -0.112646276 | 0.038142159 |
| HDLBP | 0.069442486 | 0.023878445 | GMPR | -0.104058666 | 0.023878445 |
| AKR1C3 | 0.068850177 | 0.041834939 | NDUFS3 | -0.103454163 | 0.049570705 |
| GNG2 | 0.053304921 | 0.039750765 | CDK2 | -0.101733949 | 0.041299532 |
| RRBP1 | 0.051934927 | 0.044278362 | SERPINF2 | -0.099846011 | 0.03266895 |
| UTRN | 0.05130186 | 0.041285251 | IGFALS | -0.098880263 | 0.042958346 |
| UGDH | 0.050719444 | 0.041299532 | RAB3D | -0.097122612 | 0.03098785 |
| RHOC | 0.044872949 | 0.032701836 | HBD | -0.096654064 | 0.039522413 |
| EEF1A1 | 0.044349219 | 0.049539663 | F2 | -0.092982349 | 0.03098785 |

### Adipose tissue-derived EVs from individuals with overweight or obesity promote proliferation of ER-positive breast cancer cells

To examine acute *versus* sustained effects of breast adipose tissue-derived EVs on breast cancer cell proliferation, we either performed a single treatment or long-term education of breast cancer cells prior to assessing proliferation rate. In the acute setting (Fig 2A), treatment of MCF7 cells with a single dose of 1μg/ml human adipose tissue-derived EVs showed no effect on cell proliferation over a 90-hour period, as assessed using the xCELLigence real-time cell analyzer (Fig 2B and C). In the sustained experimental setting (Fig. 2D), breast cancer cells were treated or “educated” with three doses of 1μg/ml EVs over a seven-day period. Interestingly, EVs isolated from women with overweight or obesity (O-EVs) promoted the proliferation of MCF7 cells compared to MCF7 cells not treated with EVs, while those from women with healthy weight (L-EVs) reduced proliferation (Fig 2E; p < 0.0001). Since long-term educated MCF7 cells retained their properties after passaging cells in the absence of EVs, we next sought to determine the stability of the observed effects. The proliferation rate of cells that had been further passaged and cryopreserved was assessed. The proliferation of thawed O-EV-educated cells remained significantly higher than control cells (Fig S1A; p < 0.0001 and B; p < 0.0001). These data suggest that long-term education causes sustained alterations to breast cancer cells.

To determine whether effects of BMI on breast cancer cell proliferation were generalizable, we performed long-term education of MCF7 cells with EVs isolated from 9 lean cases and 24 overweight/obese cases. We observed that O-EV-educated MCF7 cells showed higher average proliferation rate compared to L-EV-educated cells (Fig 2F; p = 0.0215). Similarly, we found that O-EVs also promoted the proliferation of another ER-positive breast cancer cell line, T47D (Fig S2; p = 0.0186). Collectively, these findings suggest that breast adipose tissue-derived EVs from individuals with overweight/obesity promote the proliferation of ER-positive breast cancer cells.

### Mammary fat pad EVs enhance tumor growth in an allograft model

Our human findings were then modeled in mice. Four-week-old female C57BL/6 mice were ovariectomized to predispose to diet-induced weight gain and randomized into two groups, receiving either low-fat diet (LFD) or high-fat diet (HFD) for 10 weeks (Fig 2G). The body weights of mice fed a HFD were significantly higher than LFD-fed mice after 10 weeks (Fig 2H; p < 0.0001). Mammary fat pads from each group were isolated and pooled and EVs were isolated from conditioned media as described for human studies. The EO771 mammary cancer cell line, syngeneic to C57BL/6 mice, were educated with EVs isolated from mammary fat pads of the LFD and HFD groups and allografted into the mammary fat pad of chow-fed C57BL/6 mice (Fig. 2I). Tumor growth was monitored over a 4-week period. EO771 cells educated with EVs from HFD-fed mice formed tumors tended to grow faster than EO771 cells educated with EVs from LFD-fed mice (Fig 2J). One mouse (10%) in the LFD group and four mice (40%) in the HFD group died during the course of the study (Fig 2K, p = 0.1089). Tumors from these mice were not collected. At endpoint, all remaining mice in the HFD group had tumors, while 1 mouse in the LFD group was tumor-free. By study endpoint (4 weeks after tumor cell injection), there was a trend for tumors in the HFD group to be larger than those isolated from mice in the LFD group (Mean ± SEM, 0.68 ± 0.12 vs 0.42 ±0.14, p = 0.0879) (Fig. 2L).

### O-EVs induce proliferation of breast cancer cells by enhancing mitochondrial respiration

To decipher the mechanism mediating the increased proliferation caused by O-EVs, we performed RNA-Seq on O-EV-educated MCF7 cells compared with control cells. Ingenuity Pathway Analysis (IPA) revealed several pathways that were significantly altered in the O-EVs educated cells, including oxidative phosphorylation (OXPHOS), EIF2 signaling, mitochondrial dysfunction, mTOR signaling and regulation of eIF4 and P70S6K signaling (Fig. 3A). The most upregulated pathway by O-EVs was OXPHOS and the genes encoding components of respiratory complexes (RC) I, III, IV and V were significantly over-expressed (Table 2). Thus, to identify the effect of O-EVs on RC function, the Seahorse XF Cell Mito Stress Test Kit was used to assess mitochondrial respiration (Fig. 3B). Basal oxygen consumption rate (OCR) of O-EV-educated MCF7 cells was significantly higher than that of control cells (Fig. 3C; p < 0.0001). ATP production by mitochondria of Ob-EV-treated cells was double that of control cells (Fig. 3D; p < 0.0001), while the maximal respiratory capacity of O-EV-educated cells was found to be significantly higher than control cells (Fig. 3E; p < 0.0001). Results were consistent across five overweight/obese cases. We also verified this observation in T47D cells (Fig. S3).

**Figure 3.**
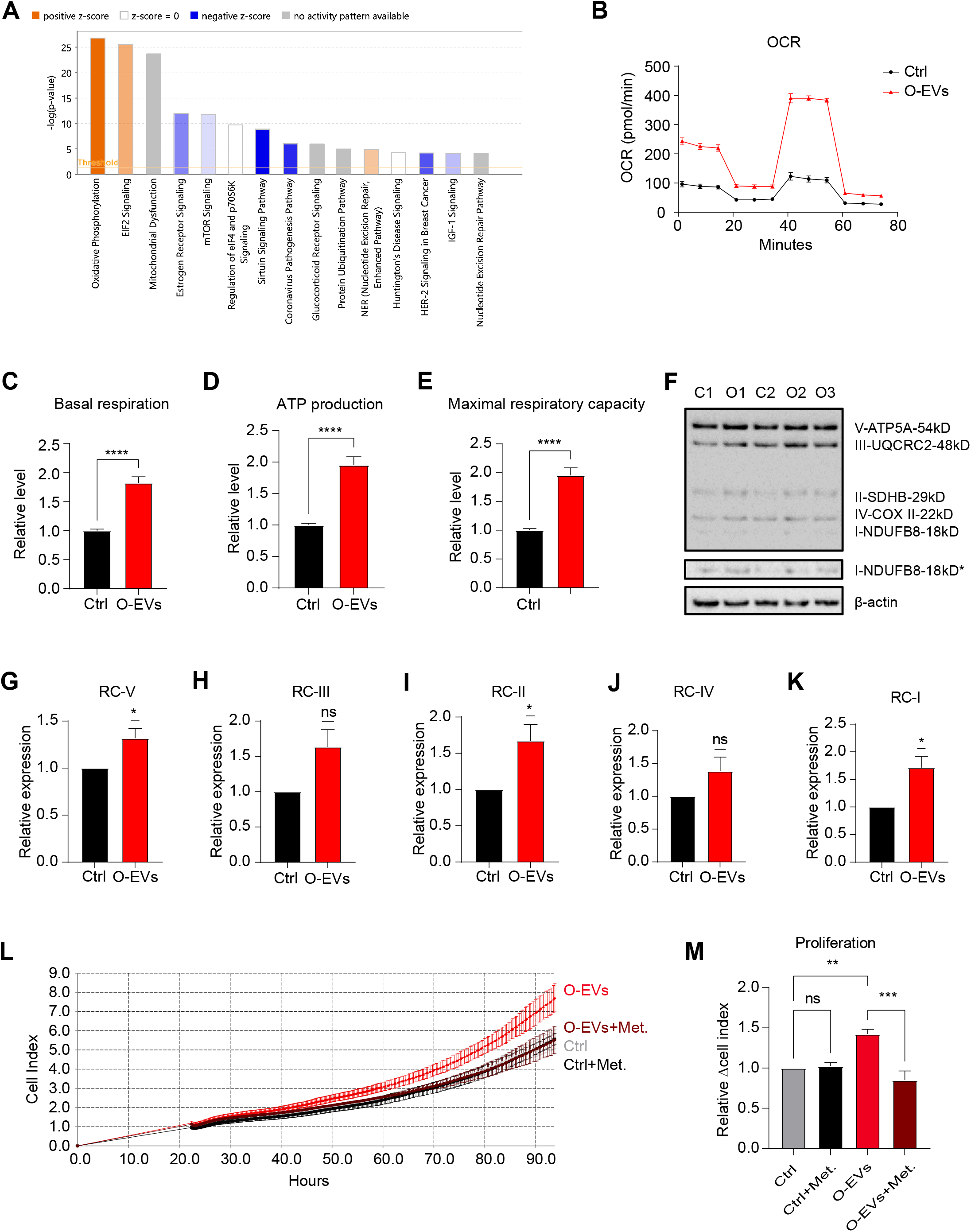
O-EVs induce proliferation of breast cancer cells by enhancing mitochondrial respiration. A) IPA analysis of RNA-Seq data revealing the top 15 pathways altered in O-EV-educated MCF7 cells compared to control. B) Representative graph of Seahorse oxygen consumption rate (OCR) of O-EV-educated MCF7 cells compared to control. (C-E) Quantification of (C) basal respiration, (D) ATP production and (E) maximal respiratory capacity of MCF7 cells educated with O-EVs from 5 independent obese cases compared to control. F) Western blot showing the levels of respiratory complexes (RCs) in whole cell extracts of O-EV-educated MCF7 cells from 3 obese cases relative to their control. β-actin was used as loading control. *The same membrane was exposed longer to show clearer bands of RC-I. (G-K) Band intensity of (G) RC-V, (H) RC-III, (I) RC-II, (J) RC-IV and (K) RC-I was quantified from western blots of 5 obese cases and normalized to control. L) Representative xCELLigence data from metformin-treated O-EV-educated MFC7 cells. M) Analysis of effects of metformin on cell proliferation across 3 independent cases. All proliferation rates are normalized to non-metformin-treatment controls. Bars and error bars represent means ± SEM; statistically significant by unpaired t-test with Welch’s correction (C-E), One-sample t test (G-K) and one-way ANOVA (M), (ns P≥0.05, *P <0.05, **P <0.01, ***P <0.001, ****P <0.0001).

**Table 2.**
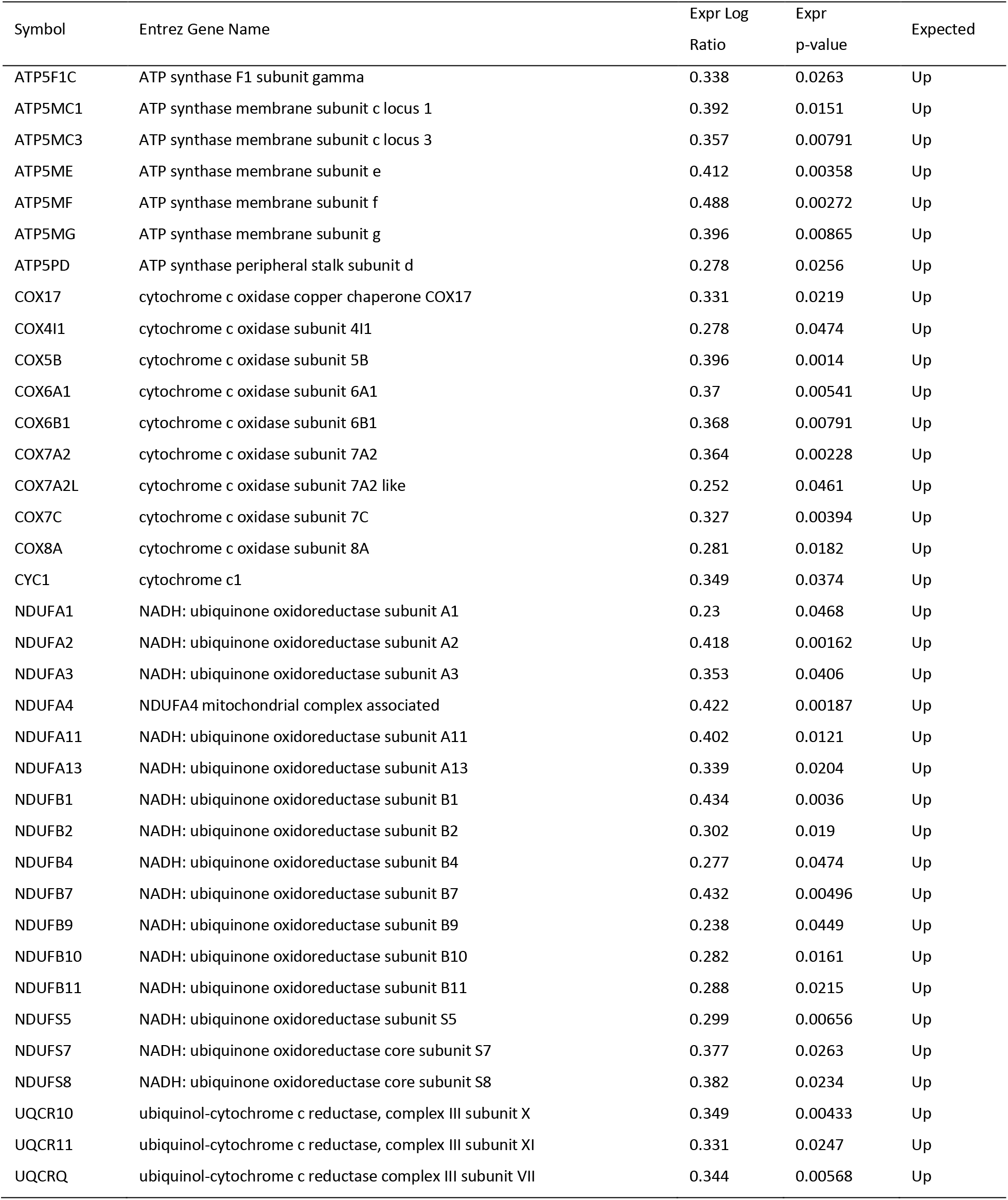
OXPHOS genes upregulated in response to long-term education with O-EVs

Consistent with Seahorse data, western blotting showed that protein levels of RC I, II, III, IV and V were higher in O-EV-educated MCF7 cells compared to control (Fig. 3F), with RC-V (p = 0.0361), RC-II (p = 0.0407) and RC-I (p = 0.0240) being significantly higher after quantification (Fig. 3G-K).

To further confirm whether the stimulated proliferation in the O-EVs educated MCF7 cells was attributable to elevated OXPHOS, cells were treated with metformin, known to inhibit mitochondrial complex I and ATP synthesis ^34^. Metformin treatment dramatically inhibited proliferation of O-EV-educated cells, while having no effect on control cells (Fig. 3L), indicating the increased dependence of O-EV-educated cells on mitochondrial respiration. This phenomenon was consistent when the assay was repeated with cells educated with EVs from three different overweight/obese cases (Fig. 3M).

### O-EVs increase mitochondrial density

To determine whether O-EVs stimulated mitochondrial respiration in breast cancer cells via effects on mitochondrial density, mitochondria were stained with MitoTracker Green FM (MTG). MTG is a mitochondrial-selective fluorescent label used to assess mitochondrial distribution and density, without membrane potential involvement. O-EV-educated MCF7 cells exhibited higher MTG intensity per cell (Fig. 4A). Data based on cells educated with O-EVs from 3 cases demonstrate that MTG is significantly higher in O-EV-educated cells *versus* control (Fig. 4B; p < 0.0001).

**Figure 4.**
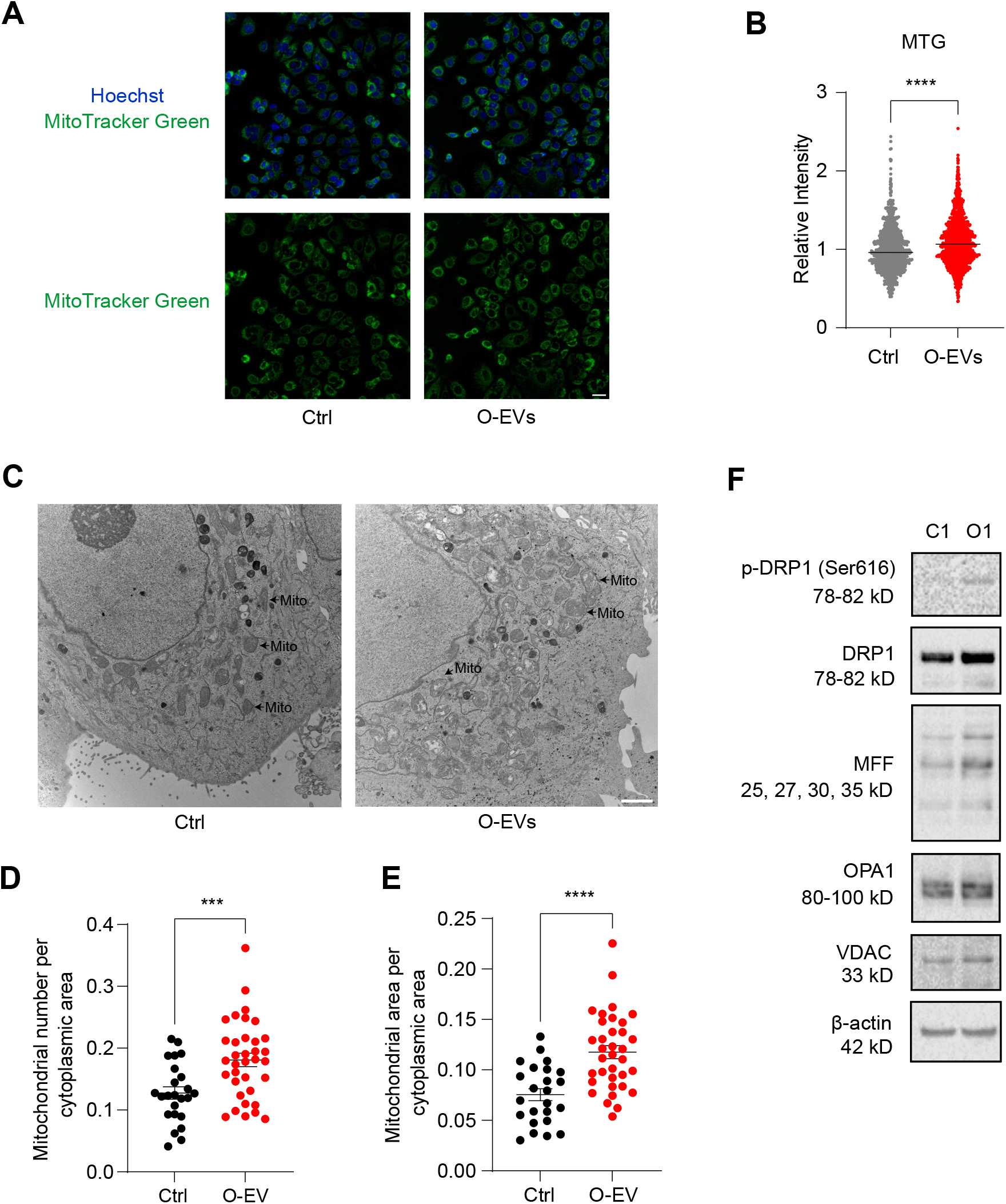
O-EVs increase mitochondrial density. A) Representative image of MitoTrakcer Green (MTG) fluorescent staining of MCF7 cells educated with O-EVs or control. Scale bar, 30 μm. B) Quantitative analysis of mean MTG intensity based on 3 obese cases vs control. MTG intensity was quantified using Imaris. C) Representative transmission electron microscopy (TEM) images of mitochondrial distribution in O-EV-educated cells compared with control. Arrows indicate examples of mitochondria. Scale bar, 2 μm. D) Mitochondrial density of O-EV-educated MCF7 cells compared to controls based on TEM images. Mitochondrial density was determined by dividing mitochondrial number by cytoplasmic area. Data are from 3 obese cases compared to control (12 fields per case). D) Mitochondrial area per cytoplasmic area of O-EV-educated MCF7 cells vs control. Data were from 3 obese cases (12 fields per case). (E) Western blot showing levels of proteins related to mitochondrial fission (DRP1, MFF) and fusion (OPA1), and mitochondrial marker VDAC. β-actin was used as loading control. Experiments were performed on MCF7 cells educated with EVs from 3 independent obese cases and 1 pair was presented. Bars and error bars represent means ± SEM; statistically significant by Mann-Whitney test (B), unpaired t-test with Welch’s correction (D-E), (***P <0.001, ****P <0.0001).

Transmission electron microscopy (TEM) was then performed on MCF7 cells educated with EVs from 3 overweight/obese cases (Fig. 4C). Quantification of mitochondria per cytoplasmic area revealed that O-EV-educated cells had a significantly higher mitochondrial density relative to control cells (Fig. 4D; p = 0.0005). By comparing mitochondrial area per cytoplasmic area, we found that O-EV-educated MCF7 cells possessed higher mitochondrial area (Fig. 4E; p < 0.0001).

Mitochondria are in a highly dynamic state between fission and fusion ^35^. In order to explore whether mitochondrial fission and fusion contribute to mitochondrial number alterations, we identified levels of proteins relevant to mitochondrial fission and fusion by WB (Fig. 4F). Phosphorylation and expression of DRP1, a key factor involved in the initial stage of mitochondrial fission, were elevated in O-EV-educated cells. The adaptor protein, mitochondrial fission factor (MFF), which is as an indicator of mitochondrial midzone division ^36^, was up-regulated in the O-EVs-educated cells, while fusion-related protein OPA1 was not affected.

### miRNAs mimic some of the effects of O-EVs

Given that miRNAs have been reported to mediate some effects of EVs ^12,37,38^, we identified several miRNAs that were enriched in O-EVs relative to L-EVs (Table 3) which could account for gene expression changes observed in O-EV-educated MCF7 cells (Table 4). Common miRNAs included hsa-let-7a-5p, hsa-miR-155-5p, hsa-miR-10a-3p, hsa-miR-29b-3p, hsa-miR-30a-3p and hsa-miR-30b-5p. T47D cells were transfected with miRNA mimics using a long-term education strategy, similar to education with EVs, and collected for xCELLigence proliferation assays, and Seahorse assay. hsa-miR-155-5p, hsa-miR-10a-3p and hsa-miR-29b-3p significantly enhanced mitochondrial respiration, including basal respiration, ATP production, and maximal respiratory capacity (Fig 5A-C, p ≤ 0.05). hsa-miR-155-5p and hsa-miR-30a-3p significantly stimulated breast cancer cell proliferation, while hsa-let-7a-5p, hsa-miR-29b-3p and hsa-miR-30b-5p caused a significant reduction of cell proliferation (Fig 5D; p ≤ 0.05).

**Table 3.** miRNAs enriched in O-EVs relative to L-EVs.

| miRNA | Base Mean | log2 Fold<br>Change | P value |
| --- | --- | --- | --- |
| hsa-mir-30a-3p | 55.49732 | 7.366509 | 0.000958 |
| hsa-mir-342-3p | 53.57631 | 7.328832 | 0.000707 |
| hsa-let-7a-5p | 52.08759 | 7.273319 | 0.001157 |
| hsa-mir-487b | 50.79617 | 7.241734 | 0.001072 |
| hsa-mir-30b-5p | 48.19151 | 7.170199 | 0.001073 |
| hsa-mir-125b-3p | 44.10294 | 7.040289 | 0.00139 |
| hsa-mir-204-5p | 44.53472 | 6.952919 | 0.017813 |
| hsa-mir-370 | 44.66252 | 6.947392 | 0.022114 |
| hsa-mir-381 | 37.99519 | 6.820632 | 0.002202 |
| hsa-mir-320a | 36.99797 | 6.675907 | 0.027862 |
| hsa-mir-205-5p | 36.58729 | 6.661963 | 0.026897 |
| hsa-mir-24-2-5p | 32.76049 | 6.505976 | 0.02909 |
| hsa-mir-155-5p | 31.90339 | 6.462966 | 0.033267 |
| hsa-mir-29a-3p | 31.75665 | 6.454281 | 0.034289 |
| hsa-mir-324-5p | 28.51078 | 6.382179 | 0.006875 |
| hsa-mir-148a-5p | 28.65046 | 6.377471 | 0.008738 |
| hsa-mir-106 | 27.92012 | 6.362008 | 0.005751 |
| hsa-mir-26a-5p | 27.29812 | 6.329712 | 0.00598 |
| hsa-mir-323a-3p | 27.19862 | 6.323342 | 0.006133 |
| hsa-mir-27a-5p | 28.205 | 6.288329 | 0.035954 |
| hsa-let-7b-5p | 25.92011 | 6.229799 | 0.011144 |
| hsa-mir-125b-3p | 25.81189 | 6.151068 | 0.047147 |
| hsa-mir-101-3p | 25.51869 | 6.135071 | 0.048256 |
| hsa-mir-342-3p | 25.51869 | 6.135071 | 0.048256 |
| hsa-mir-191-5p | 25.39828 | 6.131516 | 0.044737 |
| hsa-mir-889 | 24.92333 | 6.106233 | 0.044467 |
| hsa-mir-10a-3p | 24.93838 | 6.104442 | 0.046257 |
| hsa-mir-421 | 22.5079 | 6.038928 | 0.010976 |
| hsa-mir-15a-5p | 21.99081 | 5.999304 | 0.012808 |
| hsa-mir-195-3p | 19.12749 | 5.789966 | 0.019166 |
| hsa-mir-139-3p | 18.19213 | 5.718235 | 0.020274 |

**Table 4.**
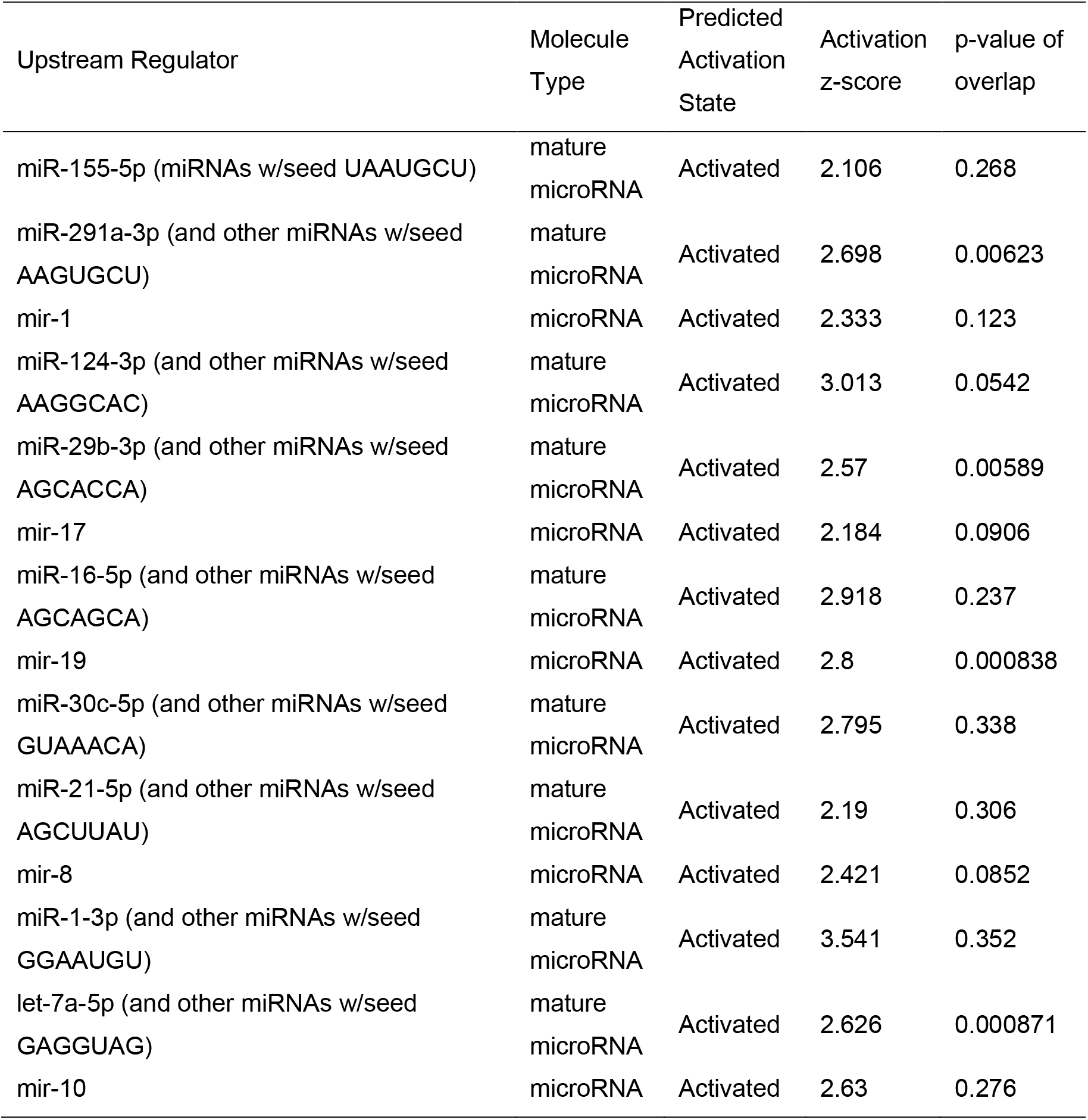
miRNAs predicted to cause observed gene expression changes in O-EV-educated MCF7 cells

**Figure 5.**
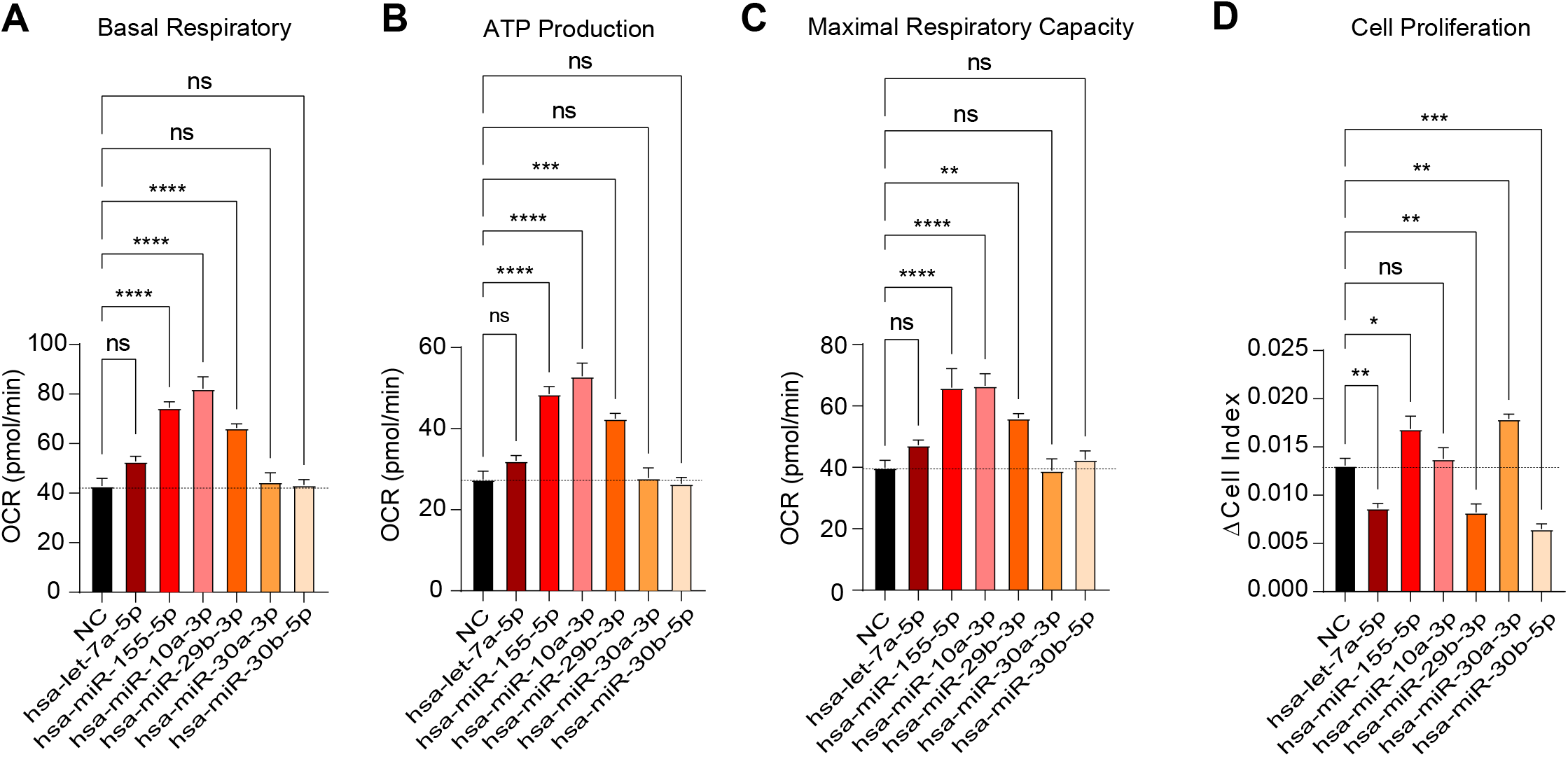
Effect of O-EV-related miRNA on OXPHOS and breast cancer cell proliferation. (A-C) Quantification of (A) basal respiration, (B) ATP production and (C) maximal respiratory capacity of T47D cells educated with miRNA mimics compared to control. D) Proliferation of T47D cells educated with miRNA mimics compared to control. Experiments were performed 4 times independently. Bars and error bars represent means ± SEM statistically significant by one-way ANOVA (A-D), (ns P≥0.05, *P <0.05, **P <0.01, ***P <0.001, ****P <0.0001).

### The effect of O-EVs on breast cancer cells is mediated by the Akt-mTOR-P70S6K signaling pathway

The Akt-mTOR-P70S6K pathway is a key regulator of both cell proliferation and mitochondrial activity ^39^. Western blot analysis demonstrates that phosphorylation (Ser473) and total Akt protein is increased in O-EV-educated MCF7 cells compared to control (Fig. 6A). Consistently, phosphorylation and total protein levels of downstream targets, P70S6K (Thr389) and 4EBP1 (Ser65) are also elevated in O-EV-educated cells versus control (Fig. 6A).

**Figure 6.**
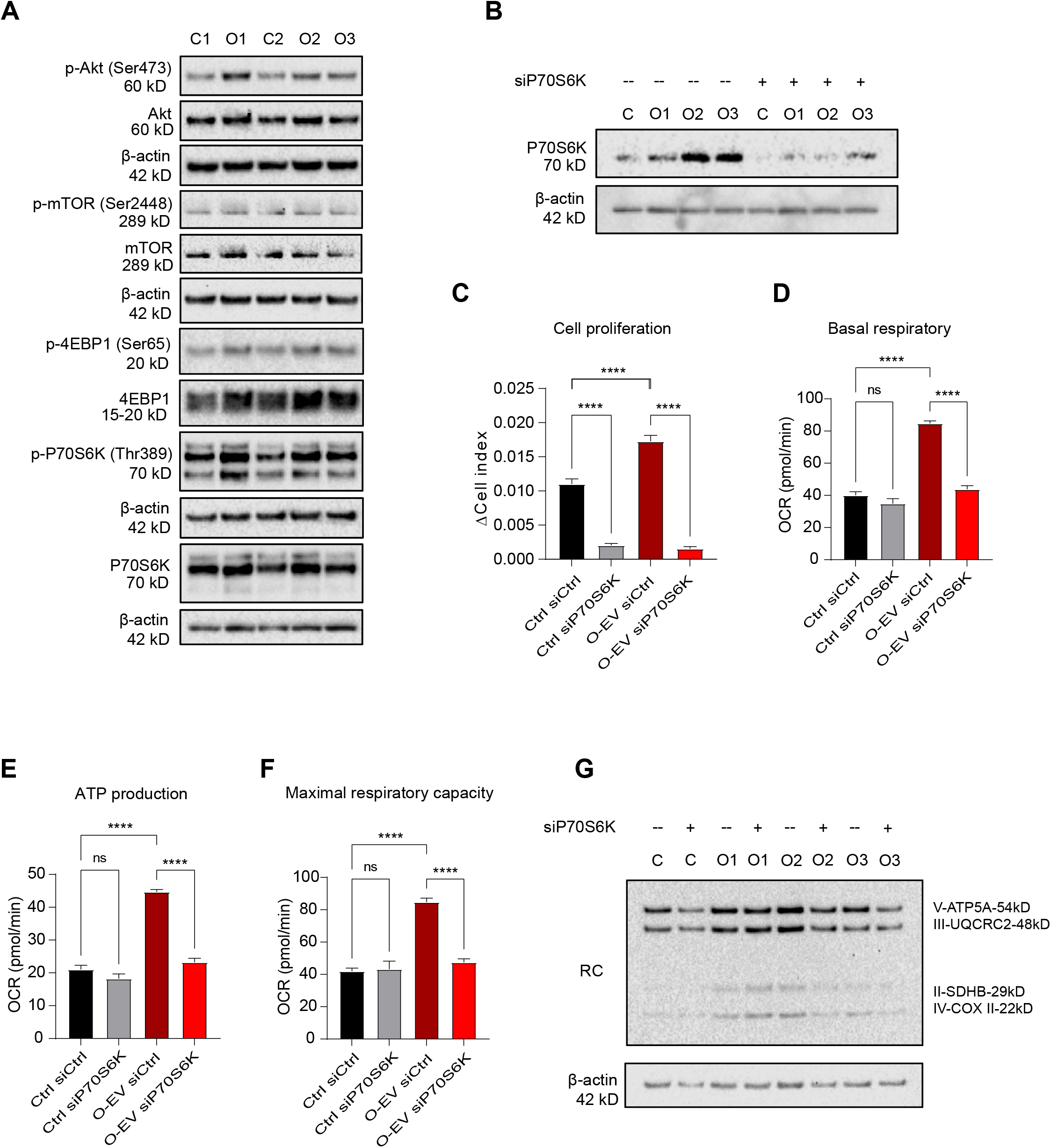
The effect of O-EVs on breast cancer cells is mediated by Akt-P70S6K signaling pathway. A) Western blot showing phosphorylation and expression levels of Akt, mTOR, P70S6K and 4EBP1 in whole cell extracts of O-EVs educated MCF7 cells (O1, O2, O3) compared to control (C1, C2). β-actin was used as loading control. Data were from 3 obese cases. B) Western blot indicating siP70S6K knockdown efficiency. β-actin was used as loading control. Data were from 3 obese cases. C) Cell proliferation assay following P70S6K knockdown. Representative experiment of 3 independent obese cases. (D-F) Quantification of effects of P70S6K knockdown on mitochondrial (D) basal respiration, (E) ATP production and (F) maximal respiratory capacity from 3 independent obese cases. G) Western blot image of the effect of P70S6K knockdown on respiratory complex expression. β-actin was used as a loading control. C: control, O: O-EV-educated cells. Bars and error bars represent means ± SEM; statistically significant by one-way ANOVA (C-F), (***P <0.001, ****P <0.0001).

To further explore whether hyperactivated P70S6K contributed to higher proliferation rates and mitochondrial respiration in O-EV-educated cells, gene silencing was performed using siRNA against P70S6K. Knockdown of P70S6K was confirmed by Western blot (Fig. 6B) and was associated with a significantly reduced proliferation of both control cells and O-EV-educated cells (Fig 6C, p ≤ 0.001). Interestingly, knockdown of P70S6K was associated with a decrease in mitochondrial basal respiration, ATP production and maximal respiratory capacity in O-EV-educated cells, but not in control cells (Fig. 6D-F). Consistently, P70S6K depletion also reduced the levels of RC proteins in O-EV-educated cells (Fig. 6G).

## Discussion

Adipose tissue dysfunction is a central mechanism driving the development metabolic disorders in individuals with obesity ^40^. This dysfunctional adipose tissue is also a key supporter of breast tumor proliferation and metastasis via altered production of inflammatory mediators, pro-angiogenic molecules and hormones, including estrogens ^11,41–43^. The interplay between adipose-derived EVs and tumor cells has been explored ^15,20,23^, but the role of EVs in driving cellular metabolic reprogramming has not been fully characterized. The strength of the current study lies in the isolation of EVs from human breast adipose tissue obtained at the time of breast reduction surgery. Given that the breast adipose tissue is complex, composed of multiple cell types that interact with each other and is significantly altered in the obese state, we opted to isolate EVs from tissue explants. This minimized the potential impact of tissue dissociation and avoided the need to differentiate cells *in vitro*, a process that does not maintain all features of obesity, and thereby provides a wholistic approach to evaluating the role of adipose tissue derived EVs on cancer cells. Long-term education was used as an approach to mimic the sustained exposure of cells to adipose-derived factors.

EVs have been extensively characterized from other sources, with their proteomic profiles being promising diagnostic tools capable of defining multiple cancers ^44^. Our study is the first to characterize EVs isolated from fresh human breast adipose tissue across a range of BMIs. We observe a trend for EVs from individuals with obesity to have a higher protein content, as previously described for melanoma ^44^. There is a significant inverse association between the number of EVs isolated per g of tissue and BMI, plausibly related to tissue from women with higher BMIs having fewer, larger adipocytes per tissue weight. EVs from human adipose tissue are enriched for exosome markers, including HSP8A, CD81, CD9 and CD63. Proteins enriched in EVs from women with obesity relative to women of a healthy weight are, for the most part, poorly characterized in the context of obesity and/or breast cancer. AKAP2 has been shown to be upregulated in ovarian cancer and promote cancer cell growth ^45^, while UCHL1 has been shown to promote cancer stemness and chemoresistance ^46,47^. Our *in vitro* and *in vivo* results point to a role for O-EVs in tumor progression. Further investigation is required to assess specific roles for protein cargoes, whether they have a role in adipose EV-cancer cell crosstalk or participate in the proliferative phenotype induced after a long-term education of ER+ human breast cancer cell lines.

A major finding of this study is the capacity of adipose tissue derived EVs, specifically from women with BMI≥ 25, to drive proliferation and metabolic changes in target breast cancer cells. Interestingly, effects on proliferation were observed after long-term education with EVs, not after a single treatment, which may explain mechanistic differences observed by other groups following a single treatment ^20,48^. Consensus related to the role and regulation of tumor cell metabolism in disease progression has evolved throughout the years. Now considered a hallmark of cancer ^49^, dysregulated cellular metabolism was initially described as a shift in energy production from mitochondrial respiration to aerobic glycolysis, otherwise known as the Warburg effect ^50–53^. However, within the last 5-10 years, emerging evidence indicates that some more aggressive cancers revert metabolism to mitochondrial respiration ^54,55^. In the current study, RNA-Seq of breast cancer cells educated with EVs from breast adipose tissue of women with obesity demonstrated an increased expression of genes involved in oxidative phosphorylation (OXPHOS). These findings are supported by functional data demonstrating increased basal and maximal respiration rates which may, in part, be due to the increased expression of respiratory complex (RC) genes. Compared with normal epithelial ductal cells, RC I, II and IV are hyperactive in human breast cancer cells ^56^, suggesting that upregulation of OXPHOS in epithelial tumor cells is a common feature of human breast cancers. Our study further demonstrates that O-EVs enhance the dependence of breast cancer cells on mitochondrial respiration

Obesity is associated with altered human adipose cell EV miRNA profiles, adding to the potential mechanisms altering cell signaling pathways in recipient cells ^57^. Interestingly, by comparing the miRNAs from obese adipose tissue derived EVs to miRNAs accountable for gene expression changes in O-EV-educated MCF7 cells, we identified miR-155-5p, miR-10a-3p and miR-30a-3p, which we found independently promote proliferation and/or oxidative phosphorylation in breast cancer cells. These miRNAs have previously been shown to regulate proliferation, Akt phosphorylation, and downstream effectors of Akt signaling, in other cancer types ^58–60^, as well as play a role in regulating cell metabolism ^61–63^. Conversely, long-term education with the Let-7a mimetic inhibited cell proliferation, consistent with previously published work ^64^, and had no effect on mitochondrial respiration. Taken together, it is likely that multiple miRNAs contribute to balanced changes in cell proliferation and mitochondrial activity that support increased energy demands of cells exposed to obese EVs.

Activation of Akt-mTOR-P70S6K signaling has previously been shown to be associated with the upregulation of RCI, II and IV of the ETC, and increased activity of RCI, to support mitochondrial respiration ^65–68^. O-EV-educated breast cancer cells display increased protein and phosphorylation levels of members of the Akt-mTOR signaling pathway. Increases in protein levels may be a direct effect of miRNAs or protein translation. Notably, silencing of P70S6K expression dramatically reduces proliferation and mitochondrial respiration of O-EV-educated cells. Mitochondrial respiration rates are also known to be impacted by changes in mitochondrial mass and mitochondrial dynamics (fission/fusion) ^53^. Compared with control cells, O-EV-educated cells displayed a greater overall mitochondrial area and number. Mitochondria are in a highly dynamic state that involves requirements for both fission and fusion ^35^. Phosphorylation of fission-related DRP1 is elevated in O-EV-educated cells, while fusion-related factor OPA1 is unchanged, suggesting a potential role of mitochondrial fission in the metabolic changes observed. Increased expression of MFF, required for healthy mitochondrial fission ^36^, suggests a need to increase mitochondrial number to support energy demands of daughter cells. Consistent with these findings, mitochondrial fission has been shown to be correlated with worse survival of breast cancer patients ^69^, and overexpression of Drp1-binding partners-MiD49/51, are known to lead to enhanced fission and proliferation, and reduced apoptosis *in vitro* ^70^. O-EV-educated cells proliferation is reduced after metformin treatment. This commonly used antidiabetic drug inhibits RC I and suppresses OXPHOS ^34^, thereby modulating mitochondrial function and confirming the importance of metabolic reprogramming in response to adipose-derived EVs. These findings highlight a potential opportunity for the clinical use of metformin in the management of breast cancer for women with obesity.

Overall, our study provides evidence for a role of EVs derived from human breast adipose tissue to regulate multiple aspects of the breast cancer cell proliferation, including changes in cellular metabolism and paving the way for further mechanistic research. EVs contribute to linking obesity to the increased risk of breast cancer and breast cancer progression

## Acknowledgments

This work was supported by the National Cancer Institute of the National Institutes of Health grant 1R01CA215797 (KAB), the Anne Moore Breast Cancer Research Fund (KAB), National Breast Cancer Foundation grant ECF-16-004 (KAB) and the Emilie Lippmann and Janice Jacobs McCarthy Research Scholar Award in Breast Cancer (KAB).

## Supplementary Figure Legends

**Figure S1.**
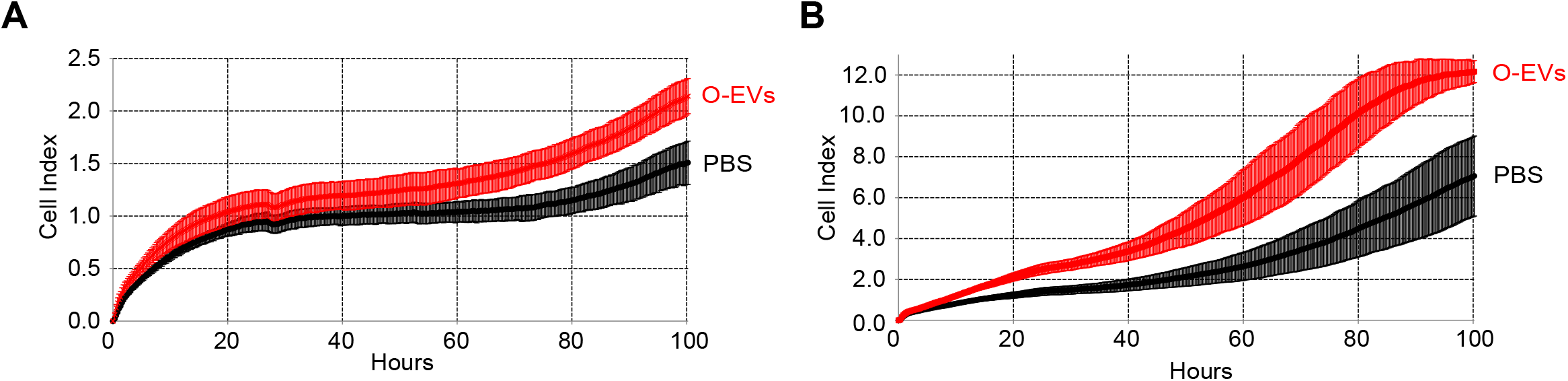
The effect of freeze-thaw on proliferation O-EV-educated cells. A) Proliferation of freshly educated MCF7 cells compared to (B) proliferation of O-EV-educated MCF7 cells after cryopreservation.

**Figure S2.**
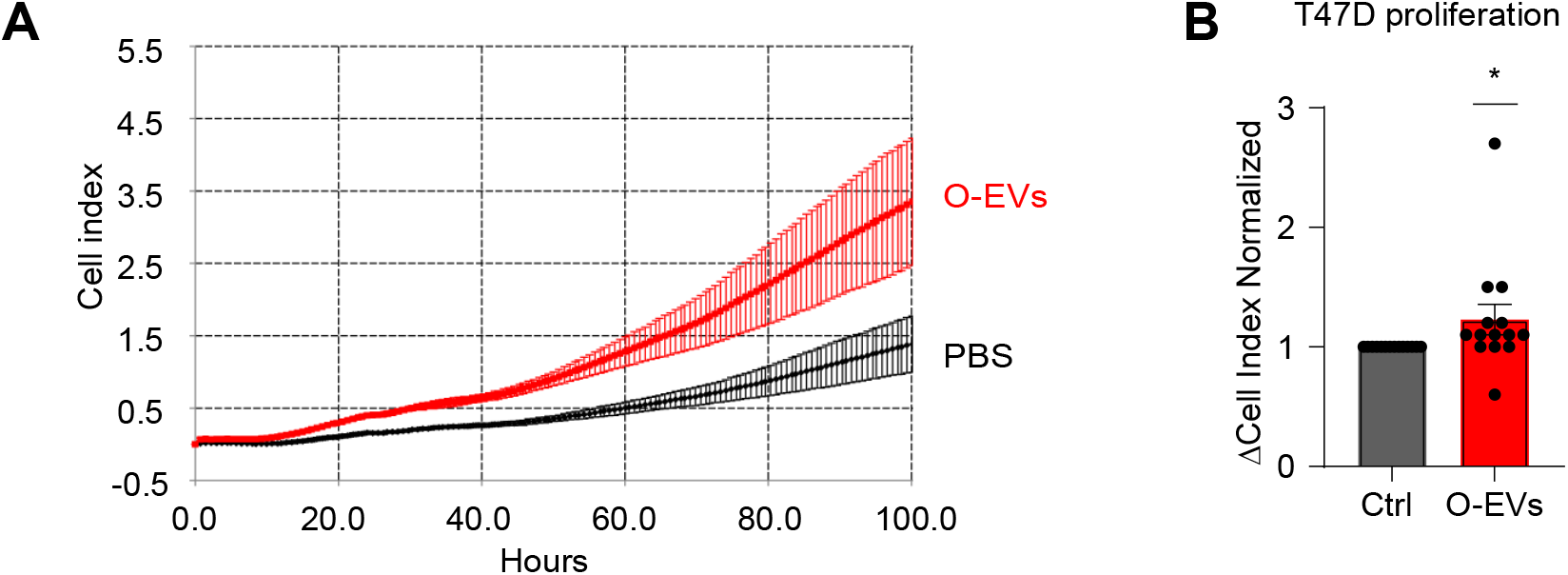
The effect of O-EVs on the proliferation of T47D cells. A) Representative xCELLigence image displaying the effect of long-term education of T47D cells with O-EVs vs control. B) Quantification of relative proliferation of MCF7 cells educated with O-EVs (n=14) vs control. Bars and error bars represent means ± SEM; statistically significant by one sample t-test (Wilcoxon test) (B), (*P <0.05).

**Figure S3.**
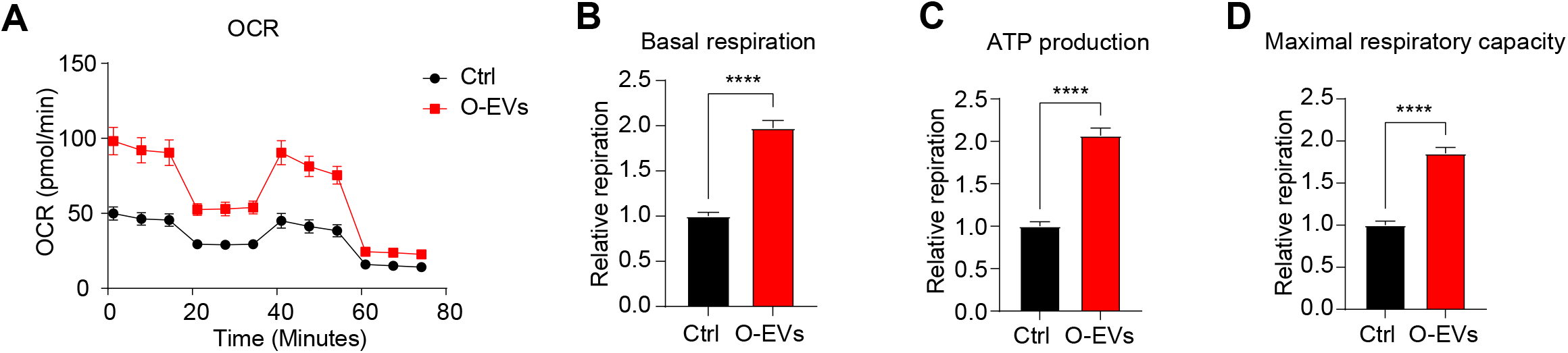
O-EVs induce mitochondrial respiration of T47D cells. A) Representative Seahorse oxygen consumption rate (OCR) image of O-EVs educated T47D cells compared to control. B) Quantification of basal respiration, C) ATP production and D) maximal respiratory capacity of T47D cells educated with O-EVs from 5 independent obese cases compared to control. Relative levels were generated by normalizing to control mean value. Bars and error bars represent means ± SEM; statistically significant by unpaired t-test with Welch’s correction (B-D), (****P <0.0001).

